# Cost-efficient boundary-free surface patterning achieves high effective-throughput of time-lapse microscopy experiments

**DOI:** 10.1101/2022.04.26.489629

**Authors:** Guohao Liang, Hong Yin, Fangyuan Ding

## Abstract

Time-lapse microscopy plays critical roles in the studies of cellular dynamics. However, to set up a time-lapse movie experiments is not only laborious but also with low output, mainly due to the cell-losing problem (i.e., cell moving out of limited field of view), especially in a long time recording. To overcome these issues, we have designed a cost-efficient way that enables cell patterning on the imaging surfaces without any physical boundaries. Using mouse embryonic stem cells as an example system, we have demonstrated that our boundary-free patterned surface solves the cell-losing problem without disturbing their cellular phenotype. Statistically, the presented system increases the effective-throughput of time-lapse microscopy experiments by order of magnitude.

## Introduction

Single-cell time-lapse microscopy has been used in various cell studies. It provides a dynamic picture of cellular regulation and has revealed many unexpected cellular regulation mechanisms [1–3] previously hidden with conventional population-based techniques. For instance, dynamic studies have helped discover factors in stem cell fate choices [4,5], the dynamics of epigenetic regulation [6,7], and the pulsatile nature of transcription in response to stress [8,9].

Time-lapse microscopy generally functions in the following way [10,11]. Live cells are placed under a microscope in a temperature, CO2, and humidity-controlled chamber to allow for long periods of cell dynamics tracking. Images are taken at a set time interval (minutes up to an hour) for an extended amount of time (hours to days) to observe the full process of interest, then all time-series images are assembled to produce a movie. If observation of multiple colonies is desired, a motorized stage could be used to move the culture plate to pre-programmed positions, aligning the objective to the desired colonies at each capturing time point. In this plate-scanning mode, images will be assembled based on their respective location in addition to time-series to produce one movie for each position, providing parallel data output.

However, this modality remains problematic for studies requiring high amplification with extended tracking time and/or large data throughput. For example, entire lineage tracking can take 4-7 days to complete [12,13] and a given colony of cells needs to be tracked from beginning to end. This is challenging because cells stay mobile throughout the tracking period and may move away from the limited field of view under high amplification, thus voiding the movie at that position. Manual adjustments to the pre-programmed positions are not feasible due to the frequency of image acquisition. Additionally, studies desiring to observe a heterogeneous population (e.g. stem cell heterogeneity [14–16]) are also hindered by this issue, since a large throughput is required to encompass the wide scenarios of a heterogeneous population. In theory, hundreds of positions on a culture dish could be pre-programmed and imaged in the plate-scanning mode, achieving large parallel cell colony input. However, cells randomly mobilizing out of the field of view means that the number of positions retaining cells at the end of recording is unpredictable. Therefore, a high effective-throughput where most if not all imaging positions retain cells throughout the time-lapse recording is key for advancing the modality.

Previous works have made many efforts to increase the effective-throughput of time-lapse movies. For instance, wide-field microscopy [17,18] has been adapted to increase the information density of time-lapse imaging. This system does not take images position-by-position, but instead takes one wide-field image (6mm X 4mm) at once. However, this system has lower resolution (700nm ~ 1.2um) [17,18], rendering it not ideal for experiments requiring high resolution to track cell divisions and movements (such as stem cell differentiation). Additionally, physical boundaries made of PDMS have also been used to confine cells for time-lapse imaging to increase the effective-throughput by minimizing cell loss in lineage experiments[1]. But such effort is not compatible with all types of cell experiment, as the physical confinement and contact induce phenotypic changes[19–21]. Furthermore, photolithography is another approach to deposit either physical or non-physical boundaries for cell confinement [22–24]. However, the high cost of this approach distances it from being widely used.

Here we propose a proof-of-principle high-informative, boundary-free, and cheap cell-patterning way to increase the effective-throughput of time-lapse microscopy. By using PDMS stencils with arrays of pre-punched holes (hundred microns diameter), we have achieved to pattern the glass surface into zones with various adhesion to cells without physical boundaries. This method confines the cultured cells to the high affinity zones at low cell density, but still allows them to grow at the low affinity zones once the high adhesive area is compact. We have demonstrated the system working for sensitive cell types (mouse embryonic stem cell) for a 4-days time-lapse movie. Simple simulation estimates that our system can at least improve the throughput by an order of magnitude.

## Material and Methods

### PDMS stencil with holes

To create isolated spots of cell adhesive surface, we have fabricated a PDMS stencil containing numerous holes (~250um diameter). First, we made PDMS using the Sylgard 184 silicone elastomer kit (10:1 base/curing ratio by weight). Second, a blank PDMS sheet of 250um thickness was manufactured by allowing liquid PDMS (degassed by vacuum half hour) to solidify in a metal mold (Fig 1A). The mold was fabricated by a Caltech machine shop (The GALCIT Shop), but it can also be ordered commercially in most off-campus machine shops. Then, the PDMS sheet was transferred onto a thin (~ 1mm thick) mylar support and placed on the bed of a VLS350 (Universal Laser Systems, Arizona, USA). To produce holes, a pattern was created using the CorelDraw software (Corel, Ottawa, Canada) with a light gray shape (color 0xFDFDFD) where a field of holes is desired. The laser then performed a raster scan at 100% power (Movie 1), and the scan was repeated three times to ensure the holes fully penetrate the PDMS membrane. Raster scanning was used because it is considerably faster than cutting small holes using the vector mode. Finally, a vector cut was performed to cut along the circular outline of the stencil (Movie 2), with a notch placed so we can determine the stencil’s orientation, as the bottom surface is much smoother and adheres better to glass than the top surface. The stencil was then washed by 95% EtOH to remove vaporized PDMS residue and sterilized.

**Fig 1.**
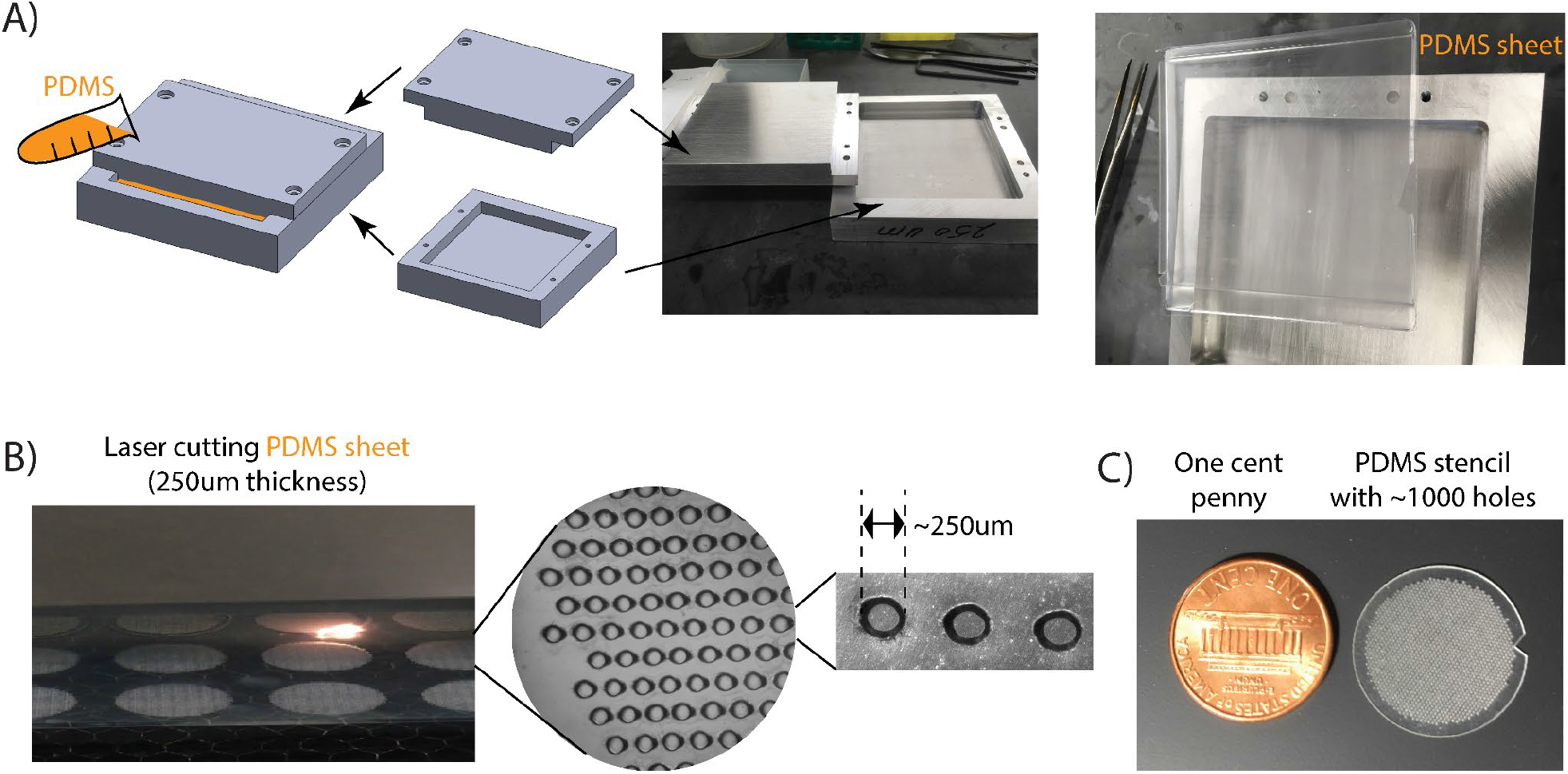
We have fabricated reusable PDMS stencils by laser cutting. (A) We used a metal mold, with a 250um gap, to manufacture a degassed PDMS sheet of 250um thickness (more details described in Materials and Methods). Worth mentioning, PDMS sheets with similar thickness are also commercially available. (B) We used a laser printer to penetrate holes (~250um diameters) on the PDMS sheet (more details in Materials and Methods). (C) About 1000 holes were created in a ~19mm diameter area, a size compatible to pattern a 35mm glass-bottom cell culture dish.

### Surface patterning

With the fabricated PDMS stencil, we have achieved to coat the glass surface into a cell adhesive pattern. First, we used 95% EtOH to clean the PDMS stencil and applied nitrogen air to blow dry. Second, we put the stencil on top of the glass bottom (Ibidi® 35mm dish 81218-200). One can improve the attachment either by manually squeezing the PDMS onto the glass surface or using a quick vacuum pressure. Third, we performed the plasma treatment, at pressure 800-1000 with medium RF level for 10 mins using the plasma cleaner (PDC-32G, HARRICK PLASMA). Fourth, we added 10ug/ml laminin (human laminin-511, BioLamina) into the dish (about 250ul, the exact volume may vary, as long as it can cover the glass surface and PDMS stencil), with a 5 mins vacuum treatment (to get rid of bubbles in the stencil holes), then incubated at 37C for 1hr. Fifth, we washed the dish with PBS three times, added 0.2% BSA-Biotin solution (Pierce™ Bovine Serum Albumin, Biotinylated Catalog number: 29130), and removed the PDMS stencil. After incubating 1hr at 37C, we washed the dish with PBS three times again. Finally, we added the labeled Streptavidin (Alexa Fluor® 405 conjugate Catalog number: S32351), incubated 10 mins at room temperature, then washed the dish again with PBS. Despite the patterned glass surface capable of being stored in 4C for days, we recommend using a freshly prepared sample for time-lapse movie experiments.

### Culture conditions and Cell line construction

Brachyury-GFP mouse embryonic stem cells (E14.1, 129/Ola, from previous publication [25], and we did not test for mycoplasm) are cultured in a humidity controlled chamber at 37°C with 5% CO2, and plated on tissue culture plates pre-coated with 0.1% gelatin (relatively cheaper compared to laminin, better option for daily culture) and cultured in standard pluripotency-maintaining conditions [26,27] using DMEM supplemented with 15% FBS (ES qualified, GIBCO), 1mM sodium pyruvate, 1unit/ml penicillin, 1μg/ml streptomycin, 2mM L-glutamine, 1X MEM non-essential amino acids, 55mM β-mercaptoethanol, and 1000Units/ml leukemia inhibitory factor (LIF). Cells were passaged by Accutase (GIBCO). For differentiation experiments, cells were cultured in the same medium, without LIF, but adding 3uM CHIR.

To label the nucleus with CFP, we have transfected a plasmid with H2B-mCerulean into the Brachyury-GFP E14 cell lines. First, we have cloned H2B-mCerulean into a pcDNA5 expression vector under a PGK promoter. Second, we have followed the standard protocol of FuGENE® HD Transfection Reagent (Promega Corporation) to perform the transfection. Then, the surviving transfected cells with strong CFP signal were sorted by Caltech FACS Facility. Finally, we plated the sorted ~271k cells, let it grow for another 4 days and froze them for further experiments and lab storage.

### Time-lapse microscopy imaging

For the 35cm glass-bottom dish, we plated about 25k cells with complete cell media to achieve a proper cell density: 1-2 cells per patterned area. The dish was manually swayed [28] to uniformly spread the cells, and left in the incubator for 2-3 hrs before imaging. Details of time-lapse microscopy setting have been described previously [29]. For each movie, about 100 stage positions were picked manually, and CFP images were acquired every 15 mins with an Olympus 60x oil objective using automated acquisition software (Metamorph, Molecular Devices, San Jose, CA), then differential interference contrast and GFP images were acquired at the beginning and the end of the movie.

## Results and Discussion

As described in the Materials and Methods and illustrated in Fig 1, we first used the laser to punch holes (about 250um diameter) on a PDMS sheet, which took less than 10 mins to punch ~1000 holes in a ~19mm diameter area (Movie 1 and Movie 2). Specifically, we used a PDMS sheet with 250um thickness. This is because thinner PDMS was too fragile to handle, while punching on a thicker PDMS created holes with cross slopes[30]. We have found that 250um is the optimal thickness capable of achieving relatively flat channel cross-sections (Fig 1B). As a comparison to the PDMS thickness, the laser cutting characteristics are not affected by the PDMS base/curing ratio. We did not observe any significant difference between the 5:1 and 10:1 mixing. Worth mentioning, despite the fact that we fabricated the 250um PDMS sheets using a home-made metal mold (Materials and Methods and Fig 1A), these sheets are also commercially available. Moreover, once fabricated, the PDMS stencil is reusable (we did not see a big difference after one year usage). Taken together, unlike most lithography patterning systems [31], which are expensive and equipment-heavy, our protocol is more accessible and cost-efficient for most biology labs.

Using the PDMS stencil with pre-punched holes, we have achieved to pattern a glass surface into various adhesion zones without physical boundaries. As shown in Fig 2A (and described in Materials and Methods), we used the PDMS stencil to coat the exposed area with extracellular matrix (we used laminin here, but our unshown data confirms other options, such as fibronectin, also work), then peeled the stencil off and incubated the pre-covered glass with blocker BSA. To observe where the boundary of high/low adhesion zones is, we used a biotin labeled version of BSA, which can be specifically bound to Streptavidin later (with Alexa405 pre-labeled). It is noticeable that, although we also used plasma treatment (Materials and Methods), it was performed after plating the PDMS, i.e., only treating the opened glass surface (i.e., hole-area) to better coat the extracellular matrix. This is the opposite of most lab-on-a-chip experiments, where the plasma treatment is applied to the entire glass surface then attaching PDMS wells for the physical boundaries[1,24]. In the latter case, the PDMS cannot be peeled off, but ours is detachable and reusable.

**Fig 2.**
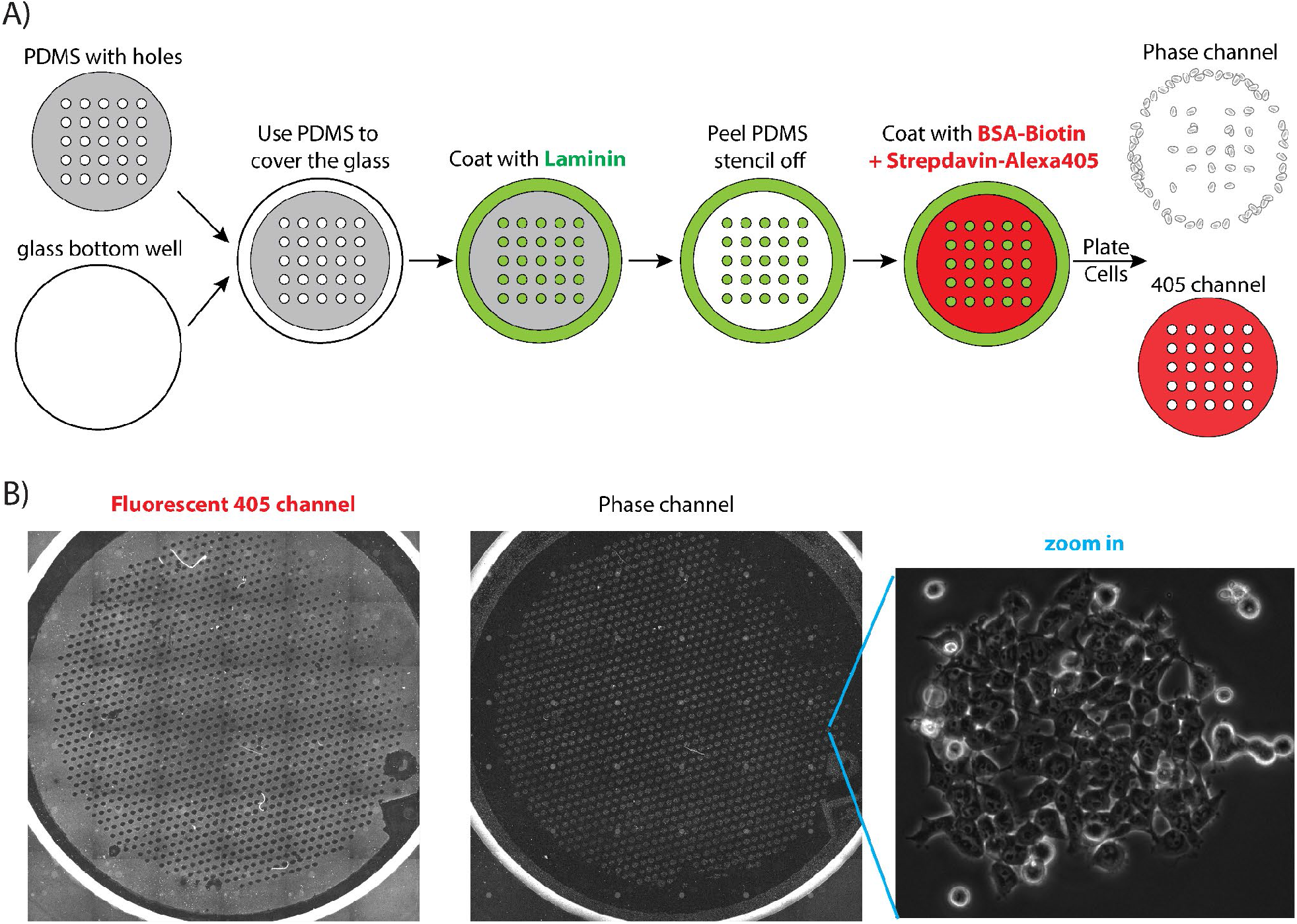
ESCs stayed in the high-adhesive patterned confined area. (A) We have used the fabricated PDMS with ~1000 holes as the stencil to coat the glass surface into high-adhesive zones (with lamin, green) and low-adhesive zones (with BSA, labeled with Alexa 405, red). (B) When plating ESCs onto the patterned surface, they formed well-shaped cell colonies in the high-adhesive zones. Worth mentioning, there are some mysterious shaded lines and colonies with high-intensity in the left and middle images. This is because, in order to cover the entire glass bottom of the 35mm culture dish, we have scanned the surface with 36 overlapping images then pieced them up. The uneven high-intensity and shade were generated due to the overlapping.

To test whether cells can be confined within the high affinity zones on the patterned surface without physical boundaries, we chose to plate mouse embryonic stem cells (ESCs). This is because: 1) This cell type is very sensitive to culturing conditions [26]. If our patterned surface works for ESC, it is more likely to be universally applicable to other cell types; 2) High effective-throughput of time-lapse microscopy (as discussed in the Introduction) is essential for ESC dynamics studies. First, high resolution imaging (using at least 60x objective) is necessary, because ESC constantly forms colonies, where cells are closed attached and difficult to segment into individual cells [26,27]. However, higher resolution often results in a smaller field of view, i.e., more serious cell-losing problems. Moreover, the high heterogeneity in the stem cell population needs, not only numerous time-lapse movies, but also the entire lineage tracking of each cell colony, to study their dynamic behaviors. Taken together, we have chosen ESC as the example system to demonstrate the efficiency of our boundary-free patterning system.

We first checked whether ESCs can sense the high-/low-adhesive zones. Specifically, we incubated a large amount of (about 250k) in a 35mm cell dish with patterned glass-bottom surface for 4 hrs, then washed the extra cells away. As shown in Fig 2B, we have found that the cells have much higher affinity to the extracellular matrix coated zone and the efficiency of the patterning is high. It is noticeable, when the surface was not covered with laminin but only patterned with BSA (such as the top right corner of the dish, Fig 2B left), no cell patterns had formed (Fig 2B middle). This observation also confirms the high specificity of ESCs’ preference on the high-adhesive zones. Worth mentioning, the no-laminin coating of the top right corner was due to a weaker attachment of the PDMS stencil edges to the glass surface. We did not further optimize the stencil attachment step, as more than 95% of the ~1000 patterned zones were forming confined patterns (Fig 2B right) and several hundreds of patterned zones are already redundant for a typical time-lapse microscope experiment. Taken together, we have found that our patterning protocol can achieve efficient boundary-free adhesion zones for ESCs.

We next tested the live cell growing circumstance, verifying whether ESCs can be restricted in the confined area in a longer term while maintaining their phenotype. Specifically, we incubated low cell density of ESCs (with 1-2 cells per confined area) at day 1, and let them grow in the standard culture condition for 4 days. As shown in Fig 3, even if the cells were initially located on the edge of the confined area, they still grew into a colony within the confined area. Admittedly, some cells did not grow into colonies, but this is because ESCs do not like culturing in a low density, regardless of the surface patterning. Remarkably, as the pattern is boundary-free, once the cell colony gets bigger, they can expand out (still centered around the confined area) (Fig 3), which is distinct from previous PDMS patterning systems with boundaries [24,31].

**Fig 3.**
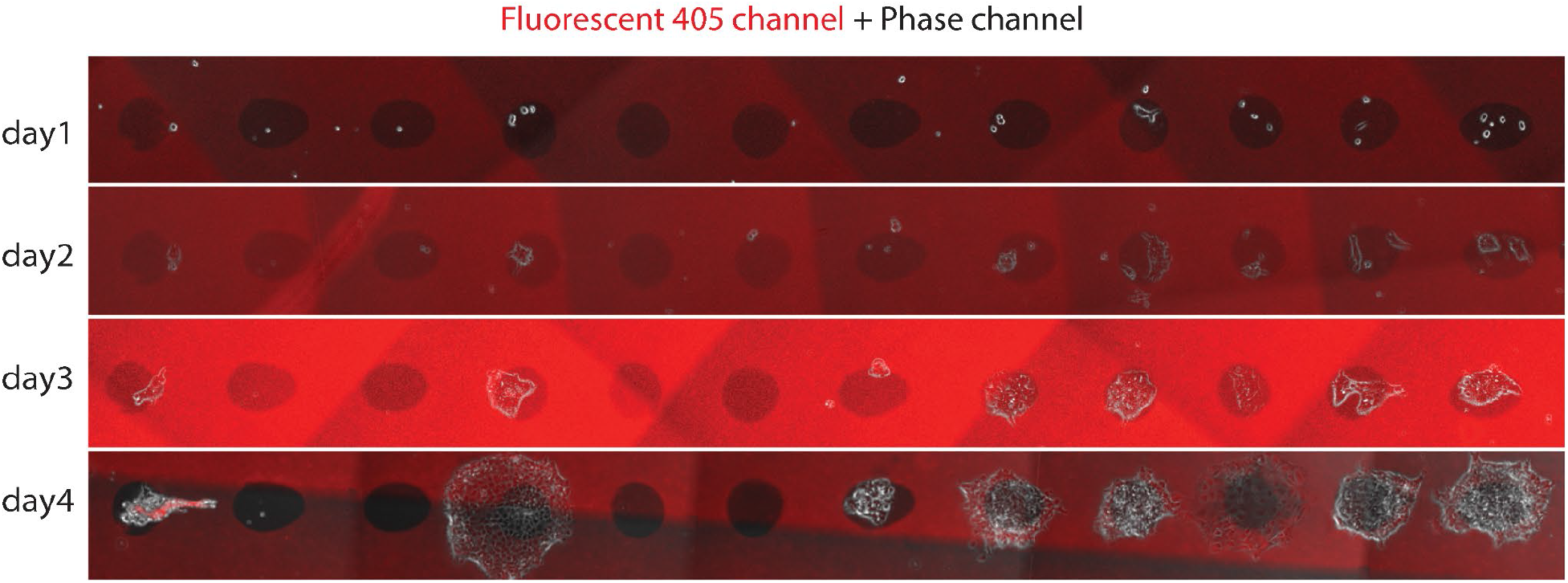
ESCs grew 4 days in the patterned surface. Cells preferred growing in the confined high-adhesive (not red) area, even if they were initially located on the edge. The pattern is boundary-free, so cells started expanding out at day 4. The red background is the fluorescent 405 channel, i.e., the low-adhesive zone. The uneven shading is due to the scanned image overlapping (similar as in Fig 2B).

We have then performed a real time-lapse movie with the patterned system. First, we fluorescently labeled the ESC nucleus with CFP (Materials and Methods, Fig 4A) and verified its pre-embedded GFP-brachyury reporter, which indicates the transition from cells’ pluripotent state to mesoderm state (data not shown). The CFP labeling, as well as the heterogeneity of CFP signals between cells, can help simplify the cell segmentation and tracking processes, as the signal is more distinguishable especially compared to the phase imaging (Fig 4B). A typical 4-days recording is shown in Fig 4C and Movie 3. Similar as in Fig 3, we have found the cells preferred growing in the high-adhesive patterned zone, and their division frequency (once every ~12 hrs) remained consistent to previous publications [32]. Once the confined area was filled, cells expanded out but still remained at a similar division time. In addition, we have observed the positive brachyury signal with high heterogeneity (as shown in Fig 4D) in 40 out 47 recorded ESC colonies at day 4, which is also consistent with previous mesoderm induction studies [32]. These results together confirm that ESCs, despite their sensitivity to culturing conditions, have mostly maintained their physiology on our boundary-free patterned surface.

**Fig 4.**
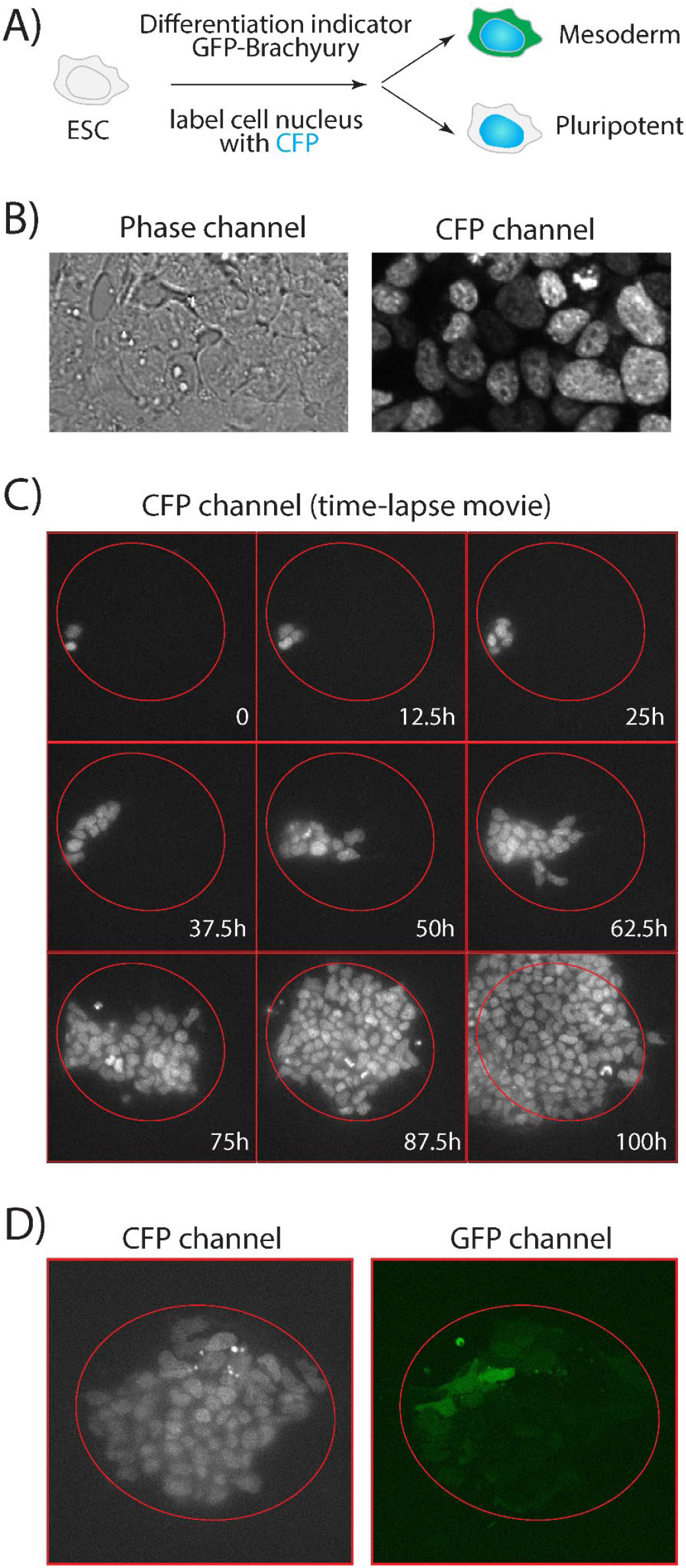
ESCs maintained their phenotype on the patterned surface in a time-lapse microscopy. A) We haved labeled the ESC nucleus with CFP. The ESC has a pre-embedded differentiation indicator GFP-Brachyury. B) CFP signal has high heterogeneity in the ESC population, which helps the cell segmentation and lineage tracking. C) 9 snapshots from Movie 3, which is a 4-days time-lapse movie of ESC on the patterned surface. The red circles indicate the confined high-adhesive area. D) A snapshot of time-lapse movie shows the ESC colony has diverse brachyury signals in the population.

Finally, we have quantified how much the patterned system can improve the effective-throughput of time-lapse microscopy experiments. First, we can track more stage positions with the patterned surface. The time interval of time-lapse movie frames is often set to be 10-20mins, because any time longer causes difficulty in cell-identification and cell-tracking due to the mobility of most cultured cells. When cells are plated randomly, the recorded positions are manually picked (a laborious process, taking hours) and mostly distance apart. As both piezo-stage moving and auto-focus processes have constrained speed, requiring about 3-10 seconds to transit from one position to another, it often takes up to half of the time in the limited time intervals (the other half is for the real imaging process). Now, with the patterned surface, both position-picking and position-transit are simplified (Fig 5A), which saves half of the transit time and enables recording ~1.5 times more positions per movie. For instance, with 15mins time-interval ESC movie tracking, we have achieved picking 119 positions in total. Second, the patterned surface solves the cell losing problem and has a significantly higher chance to record the entire dynamic lineage of an ESC colony. For instance, in a typical experiment, where we tracked 47 surviving ESC colonies in 4 days, 34 of them were recorded with a full lineage (i.e., no cell moved out of field of view, until the colony became too big and was squeezed out. By entire lineage, we define it as at least 90% of the colony were captured throughout the 4 days, like in 4C and Movie 3), and the other 13 positions captured at least half of the colonies. Together, 100% positions have captured >50% of the cell lineage and about 72.3% positions obtained the entire lineage.

**Fig 5.**
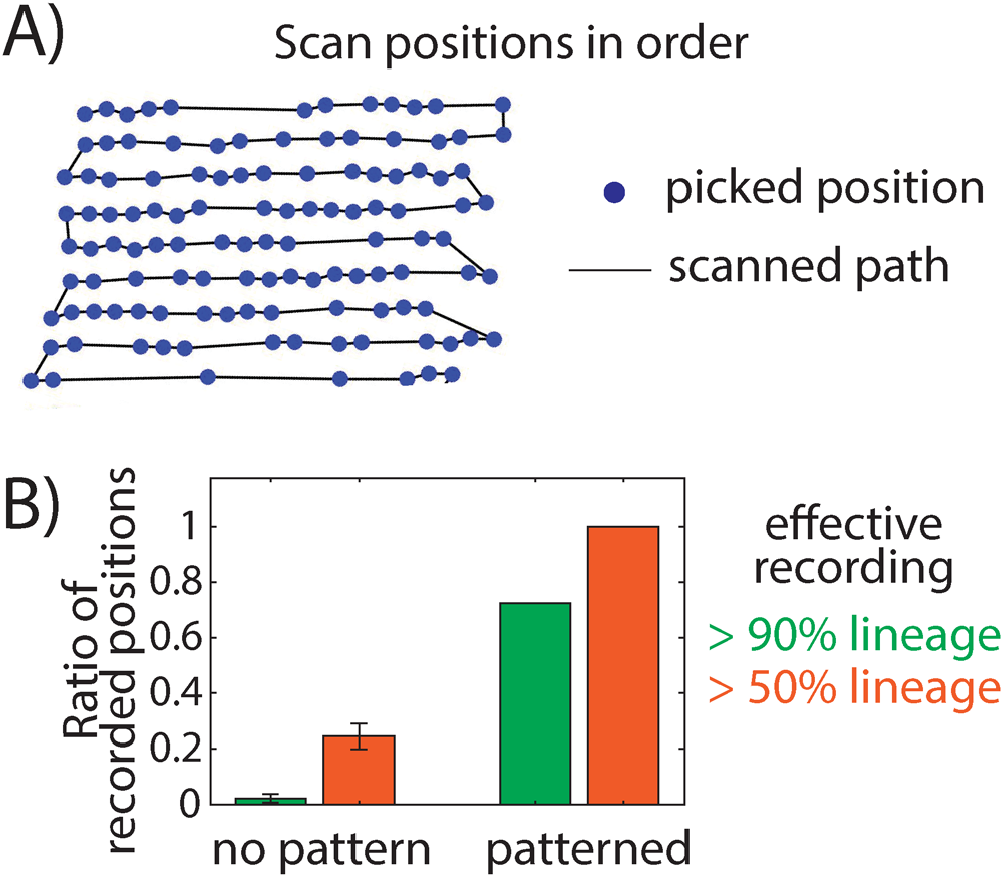
The patterned surface improves the effective-throughput of time-lapse microscopy experiments. A) Microscopy has scanned the stage positions in order. B) The majority positions of a patterned surface can record the full lineage of a cell colony, with a ratio significantly higher than the no-pattern case.

To further quantify the improvement of the effective-throughput of time-lapse microscopy experiments, we need to compare the lineage recording rate of ESCs on a patterned surface versus the no patterning case. However, without surface patterning, we can barely catch any entire lineage colony per experiment, which is impractical to collect sufficient data for the statistics. We thus performed a mathematical simulation to estimate the no-pattern situation. As shown in the Movie 4, a cell (radius 7.5um) was placed at the center of the field of view (1024 pixel by 1024 pixel, with pixel size 216.7nm, from the value of our microscopy camera). Based on our recorded movies, we set cells moving 1um per min and dividing every 12hrs (these two numbers are both consistent with previous studies too ref). Every step of a cell moving was in a random direction, and every cell division doubled cell colony size. A typical cell-losing case is in Movie 4. By simulating 100,000 movies, we found 24.5% ± 4.6% positions have captured >50% lineages, but only 2.2% ± 1.6% positions have >90% lineage (Fig 5B). Taken together, for experiments that require full lineage tracking (like ESC differentiation studies with high heterogeneity[32,33]), our patterned surface can help improve >30 times effective-throughput of a time-lapse microscopy experiment.

Worth mentioning, we have focused on culturing ESCs on the patterned surface in this work. Further optimization shall be tested for different cell types, especially about the specific choices of different blockers and extracellular matrices. For instance, we have found the protocol working less efficient for C2C12, a myoblast cell line, but mixing Pluronic F-127 with BSA can help.

## Conclusion

We have demonstrated a cheap and boundary-free surface patterning protocol that is adaptable to a wider community (especially most standard biology labs without easy access to lithography). The patterned surface has reliable bio-compatibility, even sensitive cell type ESC maintained its phenotype during a 4 days movie recording. This system significantly reduces the cell-losing problem during time-lapse microscopy movies and improves the effective-throughput per experiment by an order of magnitude. Further studies of alternative extracellular matrix and blocker options will enable this patterning protocol applicable to many other cell types and cell growth conditions.

## Author Contributions

G.L. and H.Y. contributed equally and are listed alphabetically. F.D. conceived experiments. G.L., H.Y. and F.D.. performed experiments, analyzed data, and wrote the manuscript with substantial input from all authors.

